# SMOG 2 and OpenSMOG: Extending the limits of structure-based models

**DOI:** 10.1101/2021.08.15.456423

**Authors:** Antonio B. de Oliveira, Vinícius G. Contessoto, Asem Hassan, Sandra Byju, Ailun Wang, Yang Wang, Esteban Dodero-Rojas, Udayan Mohanty, Jeffrey K. Noel, Jose N. Onuchic, Paul C. Whitford

## Abstract

Applying simulations with structure-based (Gō-like) models has proven to be an effective strategy for investigating the factors that control biomolecular dynamics. The common element of these models is that some (or all) of the intra/inter-molecular interactions are explicitly defined to stabilize an experimentally-determined structure. To facilitate the development and application of this broad class of models, we previously released the SMOG 2 software package. This suite allows one to easily customize and distribute structure-based (i.e. SMOG) models for any type of polymer-ligand system. Since its original release, user feedback has driven the implementation of numerous enhancements. Here, we describe recent extensions to the software and demonstrate the capabilities of the most recent version, SMOG v2.4. Changes include new tools that aid user-defined customization of force fields, as well as an interface with the OpenMM simulation libraries (OpenSMOG v1.0). To illustrate the utility of these advances, we present several applications of SMOG2 and OpenSMOG, which include systems with millions of atoms, long polymers and explicit ions. We also highlight how one can incorporate non-structure-based (e.g. AMBER-based) energetics to define a hybrid class of models. The representative applications include large-scale rearrangements of the SARS-CoV-2 Spike protein, the HIV-1 capsid in the presence of explicit ions, and crystallographic lattices of ribosomes and proteins. In summary, SMOG 2 and OpenSMOG provide robust support for researchers who seek to apply structure-based models to large and/or intricate biomolecular systems.

## Introduction

Over the last several decades, structure-based models have been used to study a broad range of phenomena, spanning from protein folding^1^ and binding,^2^ to the dynamics of large-scale ribonucleoprotein assemblies.^3,4^ The common feature of every structure-based model is that some (or all) of the energetic interactions explicitly stabilize specific pre-defined structures. That is, rather than assigning energetic parameters solely on the basis of chemical composition (as done in semi-empirical models), potential energy minima are defined to stabilize a known (typically experimentally-obtained) native conformation. ^5,6^ In the context of protein folding, this type of energetic representation is supported by the principle of minimal frustration,^7,8^ which states that the interactions formed in the native conformation will dominate the folding landscape. When applying these models to assemblies, such as the ribosome^9^ or protein filaments,^10,11^ assigning particular configurations to be potential energy minima can also be warranted, since structural techniques can only resolve low free-energy states. However, since structure-based models implicitly encode these free-energy minima, it is appropriate to consider the modeled interactions as representing the effective energetics of a system. With the simplicity and (generally) low computational requirements of these effective potentials, they provide a viable approach for investigating how molecular systems respond to perturbations (e.g. change in temperature, pH, binding, etc). In terms of strategies for deploying structure-based models, one will typically begin with a common-variety structure-based model,^5,6^ then systematically introduce additional energetic terms and characterize their influence on a large-scale dynamical process.

Structure-based, or Gō-like, models have a long history in the study of protein dynamics. In the earliest implementations, these models included coarse-grained (one bead per residue) descriptions of proteins, ^12^ where each residue was restricted to a 2D or 3D lattice. Subsequent models extended these coarse-grained representations to allow for off-lattice dynamics to be simulated,^5,13^ which opened avenues for direct comparison of theoretical predictions and experimental measures. The C*_α_* model by Clementi et al.5 is probably the most widely-known and used structure-based model, where countless variants have appeared in the literature over the years. In addition to the so-called single-basin models that have been popular in the study of folding, multi-basin variants^14–16^ have also been used to study conformational rearrangements. In terms of spatial resolution, all-atom^6,17^ and C*_α_*−C*_β_* ^18^ variants have been proposed to characterize side chain packing during folding, or detailed steric effects during conformational transitions. Even though all of these force fields can be categorized as being part of the same class of models, extensions and modifications have resulted in many structure-based model variants, where each has been tailored to address specific questions.

With the growing popularity of structure-based models in the 2000s, it became increasingly difficult to quantitatively compare insights provided by different model variants. Differences were typically found in the functional form and energetic weights of dihedral energies, contact maps and excluded volume terms, and different simulation protocols were often applied. A factor that drove this diversity of techniques was, at that time, there was not a generally-available tool/framework available for the application and development of new structure-based models. Rather, research groups would often design their own custom software, where idiosyncrasies would manifest in the form of seemingly-arbitrary variations that had unclear effects on the predicted dynamics.

To provide the field with a common platform for applying structure-based models, we released the SMOG server webtool (smog-server.org, or SMOG 1^19^) in 2008. SMOG 1 provided a web-based interface for generating structure-based models that could be simulated with Gromacs,^20,21^ a high-performance molecular dynamics package. With the broad support available for Gromacs-formatted force field files, some of the models generated by SMOG 1 could also be simulated in NAMD,22 LAMMPS^23^ and OpenMM.24 While the SMOG 1 interface has had considerable popularity (over 27,000 force fields generated, to date), only a few parameters could be adjusted by the user. To extend the versatility of structure-based model design, in 2015 we released SMOG 2. ^25^ SMOG 2 is distributed as a downloadable open-source software package, and it allows users to define new types of SMOG models through use of XML-based “template” files. With its lightweight and portable characteristics, users can easily disseminate SMOG-model variants without maintaining a complete software stack. Primarily driven by user feedback, the SMOG 2 package has continued to be extended and refined, and there is now a range of new options and tools for the study of arbitrarily complex biomolecular systems.

Here, we describe the most recent developments of the SMOG 2 software package, which are available in SMOG v2.4. These enhancements improve the user experience, provide new capabilities and allow for streamlined deployment of SMOG models in OpenMM (via the newly-released OpenSMOG library). To highlight the flexibility of the code, we describe example applications that test the capabilities of SMOG (v2.4) and OpenSMOG (v1.0). These examples include large systems (more than 10 million atoms, or thousands of polymer chains), very long polymers (150 kilobase strand of DNA), systems with electrostatics and explicit ions, as well as models that employ complicated functional forms that can be simulated in OpenMM. With these developments, the SMOG2/OpenSMOG tools establish a comprehensive computational infrastructure that supports the development and application of novel SMOG models using state-of-the-art hardware.

## Software Overview

The current manuscript describes two interrelated software components: SMOG (v2.4) and OpenSMOG (v1.0). SMOG 2 is a standalone software package that reads PDB-formatted structural information and produces force field files that define a structure-based model for the given biomolecular system. SMOG 2 is written in Perl, and users may install the code manually, configure the required libraries via conda, or download a fully-configured SMOG 2 container. OpenSMOG is a Python library that allows highly-customizable SMOG models to be simulated in OpenMM. While these software elements may be used independently, they have been designed to directly interface in order to expand the possible applications of SMOG models (Fig. 1). Below, we provide an overview of the support provided and usage considerations of the SMOG2/OpenSMOG framework.

**Figure 1:**
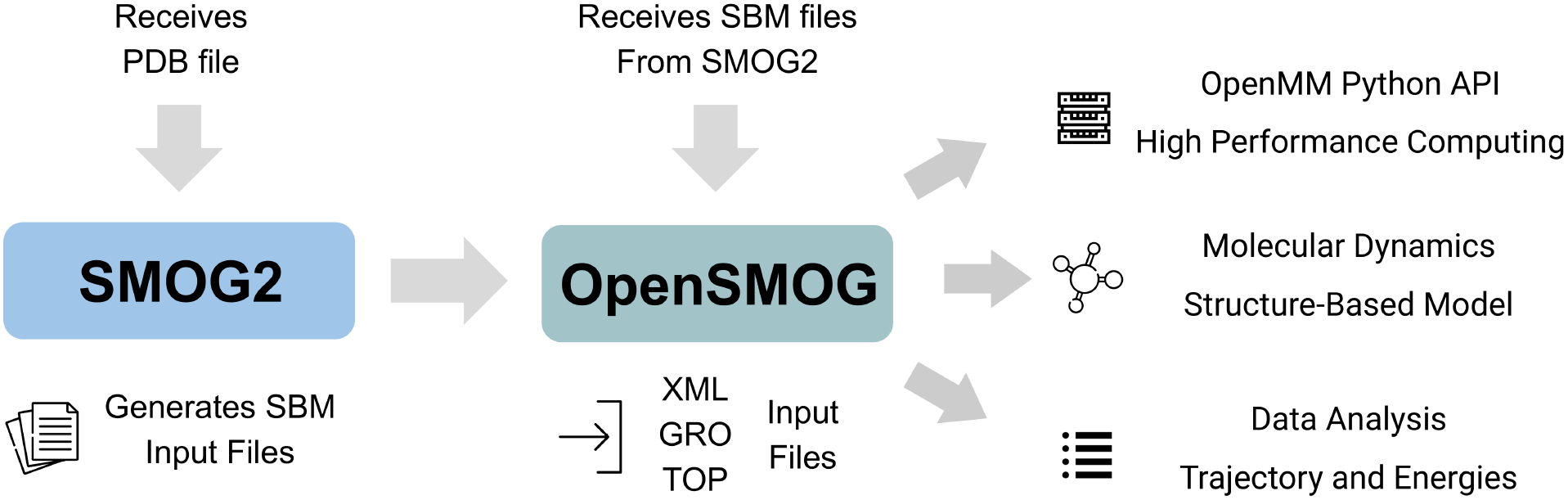
The SMOG2/OpenSMOG pipeline. SMOG v2.4 introduces support for the OpenSMOG libraries, which allow a wide range of SMOG model variants to be applied in OpenMM. While SMOG 2 and OpenSMOG are two distinct software elements, they have been co-designed to interface and provide integrated support for SMOG models. There is support for preprocessing of PDB structures, force field generation, as well as execution of GPU-accelerated simulations in OpenMM. While previous SMOG models have had general support by Gromacs, and partial support from NAMD, LAMMPS and openMM, the OpenSMOG framework extends support by enabling increased levels of model customization. In terms of work flow, SMOG 2 reads a PDB file and generates all files required to perform an MD simulation using the OpenSMOG module in OpenMM.

### SMOG v2.4

#### Approach

In addition to providing support for default C*_α_* ^5^and all-atom^6^ models, SMOG 2 allows users to define general rules for constructing new classes of structure-based models, without requiring programming or scripting. Each type of SMOG model is defined by a set of XML files, which are collectively referred to as the model “template”. These define the covalent geometry and resolution of each supported residue, the functional form of every interaction and assignment of energetics in a given SMOG model variant. There are four core files that constitute a template: the settings file (.sif), bonded information file (.bif), bond energetics file (.b) and non-bonded energetics file (.nb). SMOG 2 reads the templates, and then applies the template-defined rules to construct force fields for individual molecular systems (provided in PDB format). As output, SMOG 2 generates force field and coordinate files that can be used to perform simulations with Gromacs^20,21^ (version 3 and later) or OpenMM^24^ (via the OpenSMOG module). In addition, Gromacs-formatted files may also be used (with limited support) by NAMD,^22^ or they may be converted (through the SMOG-converter program^26^) for use with LAMMPS.^23^ Since there are already detailed technical descriptions available on template organization and SMOG 2 output files,^25,27^ we will focus the current discussion of the template-based approach to recent extensions that allow for more versatile definitions of structure-based models to be applied.

#### Code accessibility

SMOG 2 and OpenSMOG are open source and freely available for anonymous download. SMOG 2 is available at https://smog-server.org/smog2. The software may be downloaded in the form of a container, a stable release bundle, or through a git repository. Installation instructions are provided in the user manual. The OpenSMOG library can be installed via conda or pip, or it can be compiled from source (https://github.com/smog-server/OpenSMOG). Detailed installation instructions and usage tutorials can be found in the associated documentation (https://opensmog.readthedocs.io).

#### Recent improvements

##### Performance for large systems

For the current version (2.4), we made a concerted effort to improve the versatility and performance of SMOG 2 when applied to large and complex assemblies. For reference, with a typical desktop, a SMOG model can usually be generated for small molecular systems (~10,000 atoms) in less than a minute. While the performance of SMOG 2 for small systems is sufficient for routine daily usage, stress testing the code with large molecular systems revealed a number of algorithmic limitations in earlier versions. By addressing these deficiencies, we have significantly reduced the computational requirements for large systems (millions of atoms), both in terms of memory and CPU usage. For reference, as described in subsequent sections, it was not tractable to generate SMOG models for some large-scale systems (i.e. long double-stranded DNA) with SMOG v2.3. To quantify these performance gains, we compared the time required for SMOG v2.0 and v2.4 to generate an all-atom SMOG model for a reasonably large molecular system: the ribosome (~150,000 atoms). When using a 2.7GHz Intel Core i7 chip, SMOG v2.0 requires more than 8 minutes to generate force field files with the default all-atom model. Using SMOG v2.4, this same system requires just over 2 minutes. Further, as described later, the time required by SMOG 2 scales linear with the number of atoms, such that million-atom systems only require approximately 10 minutes to complete. This means it is now practical to routinely prepare structure-based model variants for larger assemblies (10^5^ − 10^6^ atoms).

##### Modeling tools

Since the release of SMOG v2.0, numerous tools (called smog-tools) have been introduced to perform common system-specific modifications to SMOG models. These modules streamline the model development stage for each user, which reduces the likelihood of introducing modeling errors and improves the reproducibility of each study. As examples, one may quickly and directly add ions to a system with the `smog_ions` tool, or rescale/remove subsets of contacts/dihedrals between specific groups of atoms (e.g. interface interactions) with `smog_scale-energies`. The `smog_extract` tool allows one to remove any subset of atoms, along with their energetic terms. This can be useful if one seeks to obtain additional sampling for a subset of atoms within a larger system. A more laborious approach to accomplish this would be to regenerate the force field files, from scratch, for a smaller system. Instead, `smog_extract` allows the user to create a force field for the subsystem that has identical energetic parameters as the complete system, including any user-introduced modifications.

##### Support for large molecules

In addition to having a range of algorithmic improvements, SMOG v2.4 can also utilize files that are provided in an extended PDB-like format. Specifically, base-62 numbering (0-9,A-Z,a-z) may be used for atom/residue indices, and free-format coordinates are allowed. With these minor changes, SMOG models can be generated for systems that have dimensions that exceed the 1-μm limit imposed by the fixed format of PDB files. With base-62 numbering, one can uniquely index and reference up to nearly one billion atoms (62^5^) or 15 million residues (62^4^) in a single chain.

##### Interface with OpenMM

While the above tools were introduced in SMOG versions 2.1 to 2.3, version 2.4 introduces the `-OpenSMOG` option, which allows for broad support of SMOG models in OpenMM. As depicted in Fig. 1, SMOG v2.4 can generate input files that are written specifically for use with the OpenSMOG module of OpenMM. In addition to enhancing support for GPU computing, the SMOG2/OpenSMOG framework also enables a more versatile approach for defining SMOG models. In SMOG 2.4, we have extended the template format, in order to exploit the Custom Force strategy implemented in OpenMM. From a user standpoint, a novel potential energy function can be added by simply introducing minor modifications to the XML-formatted SMOG2 template files. In OpenSMOG (v1.0), this framework allows for a broad range of pair-wise potentials to be deployed. Support for multi-body interactions, as well as custom nonbonded terms, dihedrals, bonds and bond angles are scheduled for future versions of SMOG2/OpenSMOG.

Here, we will show how to introduce an arbitrary contact potential in SMOG2/OpenSMOG. In the example below, we employ a custom contact potential of the form:

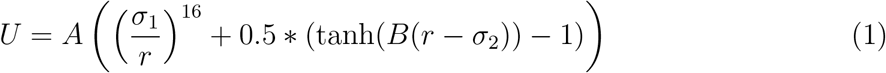

where *A*, *B*, *σ*_1_ and *σ*_2_ are parameter that define each contact interaction. While we are unaware of this specific potential being deployed previously, one could envision using this type of function to approximate a square-well interaction (Fig. 2). To define this function for use in the SMOG model, one would simply add the following element to the .sif template file:

~~~
<function name=′′r16_tanh′′
directive=′′OpenSMOG′′
OpenSMOGtype=′′contact′′
OpenSMOGpotential=′′weight*((sigma1/r)^16+0.5*(tanh(B*(r-sigma2))-1))′′
OpenSMOGparameters=′′weight,B,sigma1,sigma2′′
exclusions=′′1′′
/>
~~~

Listing 1: Defining a new contact potential in SMOG 2. By, adding this function element to the .sif file, one can introduce a user-defined potential in OpenSMOG/OpenMM.

**Figure 2:**
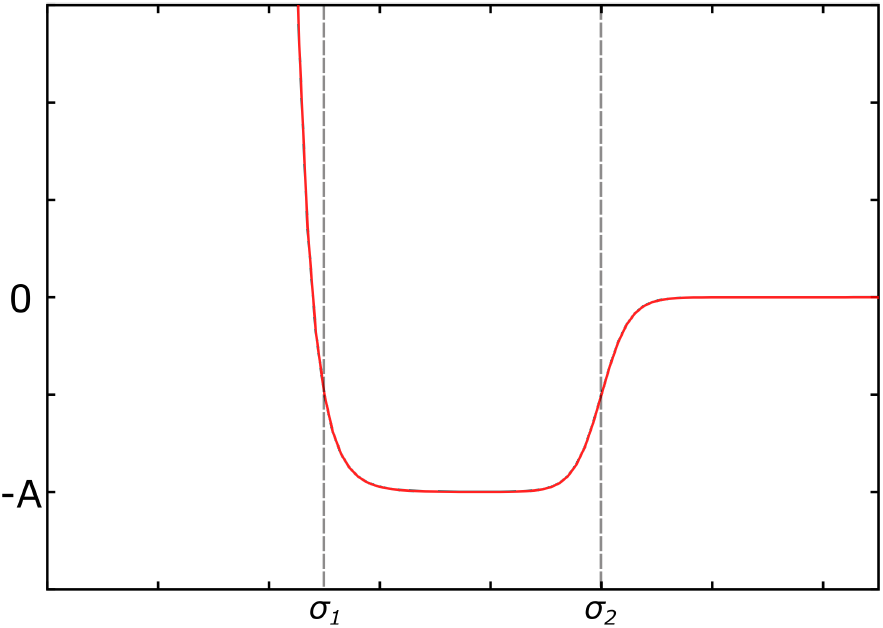
Versatile support for contact potentials in OpenSMOG. The SMOG2/OpenSMOG framework allows for a wide range of contact potentials to be employed. This figure depicts the energy function given by equation 1.

This general functional form could then be applied to any contact group, in this case the global (default) group.

~~~
<contact func=′′r16_tanh(energynorm,10,?*0.9,?*1.1)′′ contactGroup=“c“>
<pairType>*</pairType>
<pairType>*</pairType>
</contact>
~~~

Listing 2: After a contact potential is defined (Listing 1), it may be used to define the interactions between specific types of atoms in the .nb template file.

In this example, the parameter `weight` is assigned based on the default energy normalization scheme (static weights are also supported), the parameter B is 10 nm^−1^, `sigma1` is 0.9 times the native distance (calculated based on the input PDB structure) and `sigma2` is 1.1 times the native distance. After preparing the template files, the user simply needs to issue the -OpenSMOG flag when calling SMOG 2. This will ensure that all OpenSMOG-specific formatting changes are included in the produced files, so that they will be ready for use with OpenSMOG/OpenMM.

### OpenSMOG

OpenSMOG is a Python library for performing molecular dynamics (MD) simulations with structure-based models generated by SMOG (v2.4 and later. Fig. 1). OpenSMOG uses the OpenMM Python API^24^, which supports a wide variety of potential energy functions, including those used in all-atom and coarse-grained variants of SMOG models. The OpenSMOG library is a set of wrappers for core OpenMM libraries, which ensures that simulations with OpenSMOG can utilize the full range of OpenMM-supported hardware. In terms of GPU support, one may use OpenCL or HIP libraries for AMD GPUs, or CUDA libraries with NVIDIA GPUs.

Rather than developing a completely new interface for applying structure-based models, OpenSMOG builds upon existing support in OpenMM for Gromacs-formatted force field (.top) and coordinate (.gro) files. However, the utility of OpenSMOG arises from the introduction of an additional XML-formatted file with which to pass customized potential terms from SMOG 2 to OpenMM. For ease of discussion, we will refer to this file as `opensmog.xml`. This file replaces the use of the “pairs” directive in the .top file, since the pairs directive supports a limited set of pair-wise potentials. Instead, `opensmog.xml` can define pair-wise potentials that have an arbitrary number of parameters, and parameters can be set using a combination of expressions and rules, as described in detail in the SMOG 2 manual. Further, OpenSMOG passes information from `opensmog.xml` to the CustomBondForce class provided by OpenMM, which allows for use of an arbitrary number of contact potentials in a single simulation. This represents the most significant improvement over current SMOG-model support provided by other MD packages.

**Table 1:**
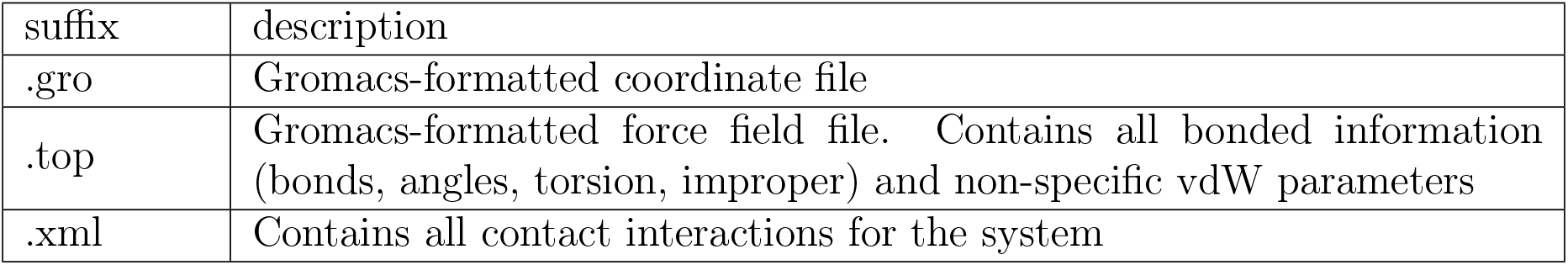
Files generated by SMOG 2 that are input for OpenSMOG.

Once the force field has been generated by SMOG 2, a short Python script is sufficient to perform a simulation with OpenSMOG. For example, to launch a simulation for the system “CI2” using HIP acceleration (for AMD GPUs), one could use the following script:

~~~
#OpenSMOG example
from OpenSMOG import SBM
sbm_CI2 = SBM(name=’CI2’, time_step=0.002, r_cutoff=1.5, collision_rate=1.0,
temperature=0.5)
sbm_CI2.setup_openmm(platform=’HIP’)
sbm_CI2.saveFolder(’output’)
sbm_CI2.loadSystem(Grofile=’CI2.gro’, Topfile=’CI2.top’, Xmlfile=’CI2.xml’) sbm_CI2.createSimulation()
sbm_CI2.createReporters(trajectory=True, energies=True, energy_components=False,
interval=1000)
sbm_CI2.run(100000, report=True, interval=1000)
~~~

Listing 3: Usage example of the OpenSMOG library

In this example, the temperature is set to 0.5 (reduced units), the simulation will be performed for 100,000 timesteps of size 0.002, where the non-bonded cutoff is 1.5 nm. There are a few important technical notes that are worth mentioning. First, a timestep of 0.002 is recommended^25^ for the default all-atom SMOG model, but a step of 0.0005 is typically appropriate for *C_α_* models. For customized models, different values may be more suitable. Next, HIP libraries are used in the above example, though OpenMM allows for OpenCL and CUDA support, and the user should determine which is most suitable for the available hardware. Finally, in the above example we set `energy_components=False`, which turns off the writing of individual energetic components (e.g. dihedrals, contacts, etc). When turned on, we have found that writing the full set of energies every 1000 steps can incur a roughly 10% performance loss. Accordingly, it is important that the user determines an appropriate frequency for writing the energies by component.

With the script given in Listing 3, OpenSMOG would generate three output files:

**Table 2:** Default output files when using OpenSMOG. Default file name is given by the variable “name” in Listing 3, with a suffix defined below.

| suffix | description |
| --- | --- |
| `_trajectory.dcd` | contains atomic positions for each frame |
| `_energy.csv` | contains the potential energy, kinetic energy, total energy and temperature |
| `_energy_components.csv` | contains the potential energy associated with each force group in the system |

## Applications of SMOG 2 and OpenSMOG

To illustrate the capabilities of SMOG v2.4 and OpenSMOG v1.0, we provide some representative examples that highlight key aspects of this computational infrastructure. First, we describe simulations of the SARS-CoV-2 spike protein, for which post-translational modifications and non-standard energetic terms are introduced and applied in OpenMM. Simulations of the HIV-1 capsid are then presented, which illustrate how one may incorporate electrostatics and explicit ions into a structure-based model. This is followed by examples of extremely large molecular systems that are supported by SMOG v2.4. These include cases that span large length scales (~5 microns), include many atoms (> 10^7^), have long chains (10^5^ residues) and systems composed of many chains (> 10^4^). These large systems were used as benchmarks for identifying performance and memory limitations in earlier versions of SMOG 2. We close with an example for how SMOG 2 may be used to easily generate a force field for subsystems of larger assemblies.

### Simulating global rearrangements with SMOG2 and OpenSMOG

SMOG2 and OpenSMOG provide a versatile framework for defining structure-based models of arbitrarily complex molecular systems. The flexibility of this approach is highlighted by recent simulations of the SARS-CoV-2 spike protein (Fig. 3). For this system, there is much interest in understanding the influence that glycans may have on the functional dynamics. For our purposes, we will consider the influence that glycan excluded volume interactions can have on the global rearrangements of this protein. That is, rather than considering the full range of energetic factors that may be present, these initial simulations aim to directly assess the steric contribution. For this type of question, one would need to define the glycans to have no preferred configuration (except those imposed by stereochemistry) and no stabilizing contacts. Further, one must be able to describe simple (N- and O -ligated) and complex (branched) glycans. In addition to these considerations, one may also wish to encode the bonded terms based on Amber^28^ parameters, rather than those found in the structural model, as described recently for simulations of a viral capsid. ^29^ In terms of the user interface, all force field parameters are defined in the templates, while the glycan bonds are given through a PDB-file extension. Using the exact same files as input for SMOG 2, one can then generate the force field files for Gromacs (default behavior), or OpenMM (by issuing the `-OpenSMOG` flag).

**Figure 3:**
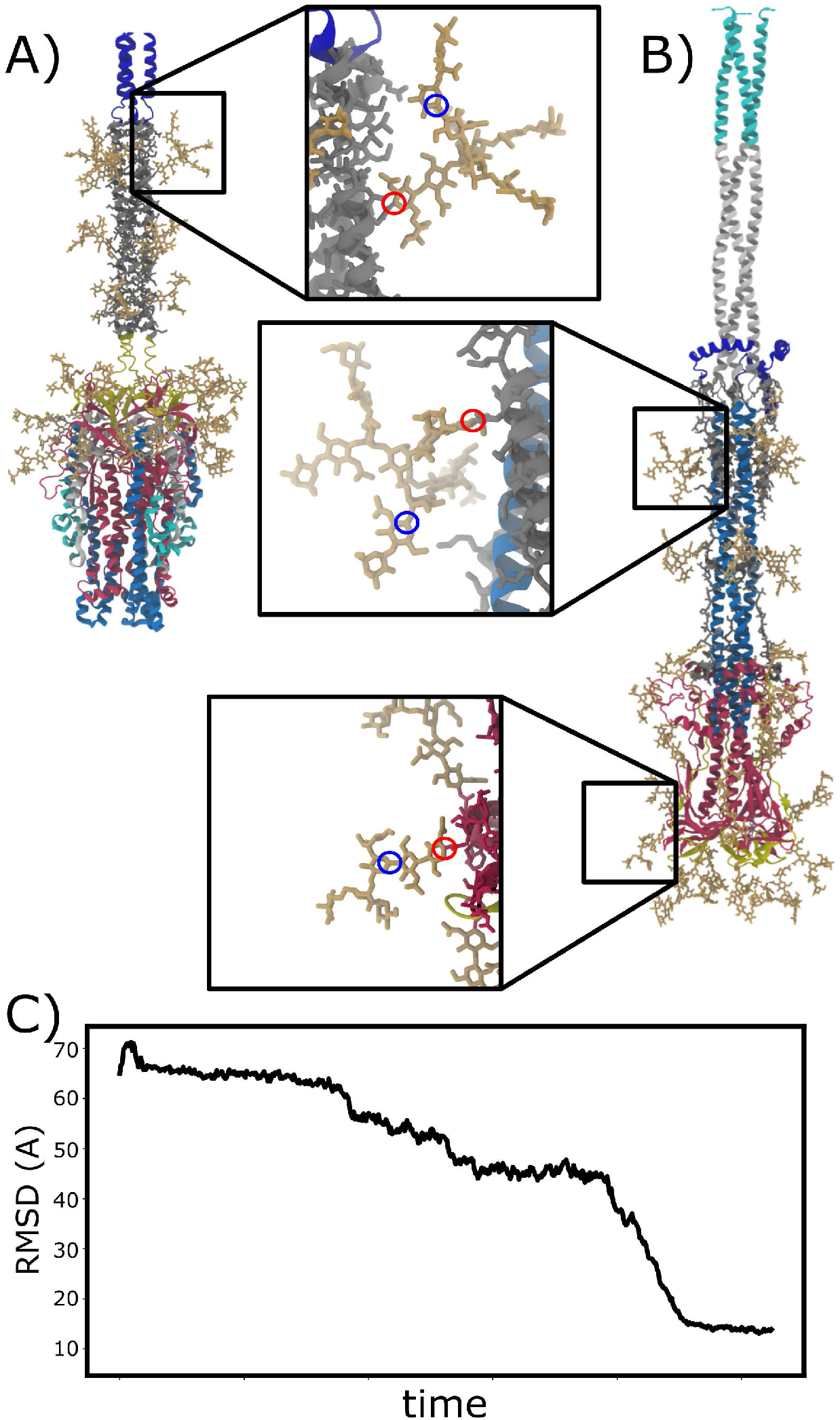
Global rearrangements of the SARS-CoV-2 Spike Glycoprotein. A) Structural model of the prefusion conformation of the S2 subunit of Spike protein. Glycans are shown in yellow. Blue circles indicate the positions of monosaccharide-monosaccharide bonds and red circles show monosaccharide-Asn bonds. B) Postfusion conformation of the S2 subunit of the Spike Protein. C) RMSD from the postfusion structure for a representative simulation with a SMOG model of the prefusion-to-postfusion rearrangement.

To demonstrate the utility of this complex definition of a structure-based model, we applied it to simulate a global conformational rearrangement of the fully glycosylated S2 subunit of SARS-CoV-2 Spike protein. During the membrane fusion process, the S2 subunit of the Spike protein undergoes a major conformational change, where individual atoms are displaced by 200-300Å. This transition is of obvious immediate interest, since it is responsible for virus-host membrane fusion, a necessary step during infection ^30–34^. However, simulating this conformational change also illustrates how complex and large-scale molecular motions may be studied using all-atom structure-based models. Following the approach described in Ref.^35^, we used this extended all-atom (15207 atoms) structure-based model to simulate the transition between the prefusion and postfusion conformations of the glycosylated S2 subunit. Here, the postfusion structure contacts are energetically stabilizing. Through use of the OpenSMOG option of SMOG 2, this complex system can be simulated using OpenMM (Fig. 3C). Alternately, one may omit this flag and perform simulations in other MD packages, such as Gromacs. In terms of computational performance, we obtained 1.73 × 10^8^ time steps per day with OpenSMOG when using a single AMD MI50 GPU.

### Studying ion dynamics in large assemblies

A goal of the SMOG class of structure-based models is to allow for the systematic analysis of factors that influence biomolecular dynamics. By starting with highly-simplified variants of structure-based models,^6^ and then continually increasing the energetic complexity, we envision that a nearly continuous spectrum of models may be constructed that span all the way to completely non-specific (e.g. AMBER, CHARMM) potentials. To this end, we continue to expand the support of SMOG 2 for more energetically detailed models. A major step in this direction is enabling one to integrate electrostatic interactions with implicit or explicit ions into the existing SMOG-model framework.

Recently, we have extended SMOG 2 to allow for effective ionic potentials to be deployed. Specifically, one may apply pairwise non-bonded interactions that include up to five Gaussian functions, as described by Savelyev and Papoian.^37^ Following their iterative refinement protocol, we established ion-protein parameters that reproduce the ionic distributions obtained with the Amber99SB-ildn^38^ force field. By developing models that have explicit diffuse ions and electrostatics, along with implicit representations of solvent, this strategy can allow for long timescale simulations of large-scale biomolecular assemblies. Since the details of this particular parameterization are described elsewhere, ^39^ we will only note the enabling technical features that are available in SMOG v2.4. First, we extended SMOG 2 in order to accommodate general non-bonded definitions. Next, we introduced the `smog_ions` tool, which allows the user to quickly assign ion positions and parameters to any existing SMOG force field. As expected, these tasks require minimal computational overhead, where assigning a million ions can be completed in a few minutes on most personal computers.

To demonstrate the potential for SMOG 2 to be used to investigate electrostatics and ionic effects in large-scale assemblies, we simulated the HIV-1 capsid with an all-atom model in the presence of 10 mM MgCl_2_ and 100 mM KCl. The HIV-1 capsid consists of 1356 p24 proteins,^36^ which together contain 2.4 million non-hydrogen atoms (Fig. 4A). Due to the dimensions of the box (cube of linear dimension 200 nm), is was necessary to introduce million diffuse ions (Mg^2+^, K^+^ and Cl*^−^*, Fig. 4B) in order to represent physiological concentrations. The simulations can be performed with a modified version of the Gromacs (v5.1.4) software package, which is available for download at smog-server.org. We then calculated the number of diffuse ions that were within 1 nm of the HIV-1 capsid, as a function of time (Fig. 4C). This demonstrative calculation shows that both monovalent and divalent ions can equilibrate around the HIV-1 capsid within a few million timesteps. This example makes it clear that it is now tractable to design new electrostatic/ion representations and investigate how these factors can influence dynamics in very large systems. While these simulations may be performed on commonly-available commodity clusters, an explicit-solvent simulation of these dimensions would contain more than 2 billion atoms, which would require specialized access and software in order to utilize world-leading computing resources^40^. In summary, these recently-introduced features of SMOG2 enable new modeling strategies to be applied in the study of ionic effects in biomolecular assemblies.

**Figure 4:**
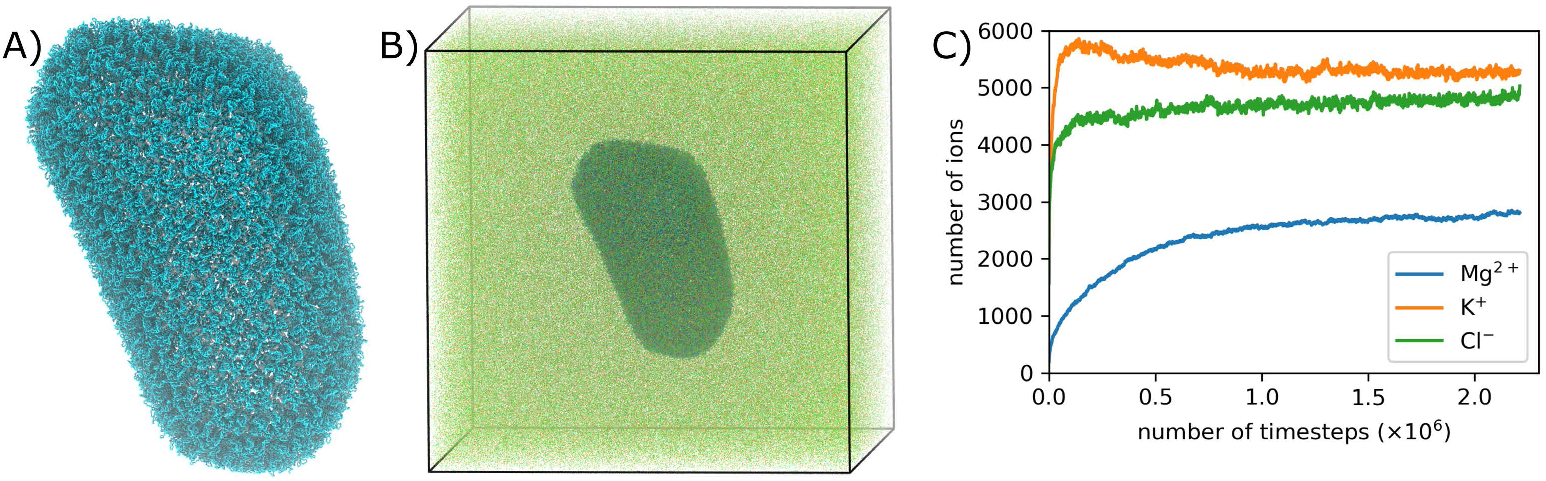
Simulations of the HIV-1 capsid with diffuse ions. A) Cryo-EM structure of the HIV-1 capsid (cyan; PDB 3J3Q^36^). B) System simulated with a SMOG model, where the HIV-1 capsid is in the presence of 10 mM MgCl_2_ and 100 mM KCl (cubic box of linear dimensions 200 nm). Diffuse ions appear as small dots. C) From a simulation with a SMOG model variant that includes electrostatics, we calculated the number of Mg^2+^ (blue), K^+^ (orange) and Cl^−^ (green) ions that are within 1 nm of the capsid. After a few million time steps, the system visibly begins to equilibrate, where the average number of each type of ion approaches an asymptotic value.

### Simulating multi-million atom systems

Since the original release of SMOG 2,^25^ a range of changes have been introduced to enhance the user experience and improve performance when working with larger molecular systems. While performing a simulation will remain the most computationally-demanding step of any study, the improved performance of SMOG 2 allows users to quickly iterate through variations of a given model, as well as troubleshoot potential issues associated with each structure and/or custom force field. To illustrate the capabilities of SMOG 2, we provide example applications to large molecular systems: crystallographic lattices of many (> 150) complete ribosomes, a lattice formed by tens of thousands of proteins and a hypothetical 1000-nucleosome system. Below we describe these benchmark systems. For the lattice of ribosomes, we also describe some basic analysis that is accessible when using SMOG models to simulate such a system.

### Crystallographic lattice of ribosomes

In order to test the support of SMOG 2 for large and heterogeneous systems, we used SMOG 2 to generate a structure-based model of a crystallographic lattice of ribosomes. In this test system, we used a crystallographic model of the T. thermophilus ribosome (PDB ID:4XEJ^41^) to first generate a complete unit cell, which is composed of eight ribosomes, for a total of 1.14 million atoms and 424 chains. Unit cell generation was performed using the Unit Cell tool in Chimera.^42^ We then generated systems with 1 unit cell (8 ribosomes) to 19 unit cells (152 ribosomes) tiled along the x direction. SMOG v2.4 could successfully generate an all-atom structure-based model (with AMBER-based bonded geometry^29^) for each of the systems using a single core of a 2.2GHz Intel Xeon Platinum 8276 CPU. The time for SMOG 2 to complete scaled linearly with the number of unit cells (~ 11.5 minutes per million atoms).

Since SMOG 2 enables one to generate all-atom structure-based models for large system, one can begin to probe how crystallographic contacts impact the apparent molecular flexibility of this large assembly. Here, we performed molecular dynamics simulation of a 15-ribosome system, which is composed of a central ribosome along with its 14 adjacent crystallographic replicas (795 chains, 2144639 atoms). After generating an all-atom structure-based model with SMOG 2, we performed constant temperature simulations (0.5 reduced units, or 60 Gromacs units) using Gromacs.^i^

Comparing the dynamics of a ribosome in isolation with the dynamics found in the lattice reveals the significant impact crystallographic contacts can have on flexibility. For this comparison, we considered the spatial root mean squared fluctuations (RMSF) of the ribosome in each system (central lattice ribosome, or isolated ribosome; Fig. 5). While many regions exhibit nearly identical RMSF values, there are distinct differences in some peripheral structural elements. In particular, there is a marked decrease in RMSF values for the small subunit beak (h35) and spur (h6), as well as the large subunit L1 stalk and protein L9. Due to the presence of crystallographic contacts, the L1 stalk, beak and spur exhibit fluctuations that are roughly linearly correlated between the lattice-bound and isolated system. This is consistent with each element moving approximately as a rigid body about a pivot-like region. That is, the mobility of each atom is roughly proportional to its distance from the pivot region. As a result, the slope of the line formed is equal to the ratio of root-meansquare-solid-angle (RMSSA) subtended by the scale of the fluctuations in the lattice/cluster and the free single ribosome. Protein L9 exhibits a different trend. That is, all atoms in L9 have roughly the same RMSF values in the lattice, consistent with strong confinement of the extended domain. In the isolated ribosome, all restraints on L9 are absent, and there is a wide range of large RMSF values. While the current simulations provide proof-of-principle that crystallographic effects may be quantitatively assessed in the SMOG 2 framework, additional calculations will be required to fully delineate their precise influence. In terms of our understanding of biomolecular dynamics, these types of comparisons provide new avenues for establishing insights into the relationship between crystallographic B-factors and dynamics of biomolecules in solution.

**Figure 5:**
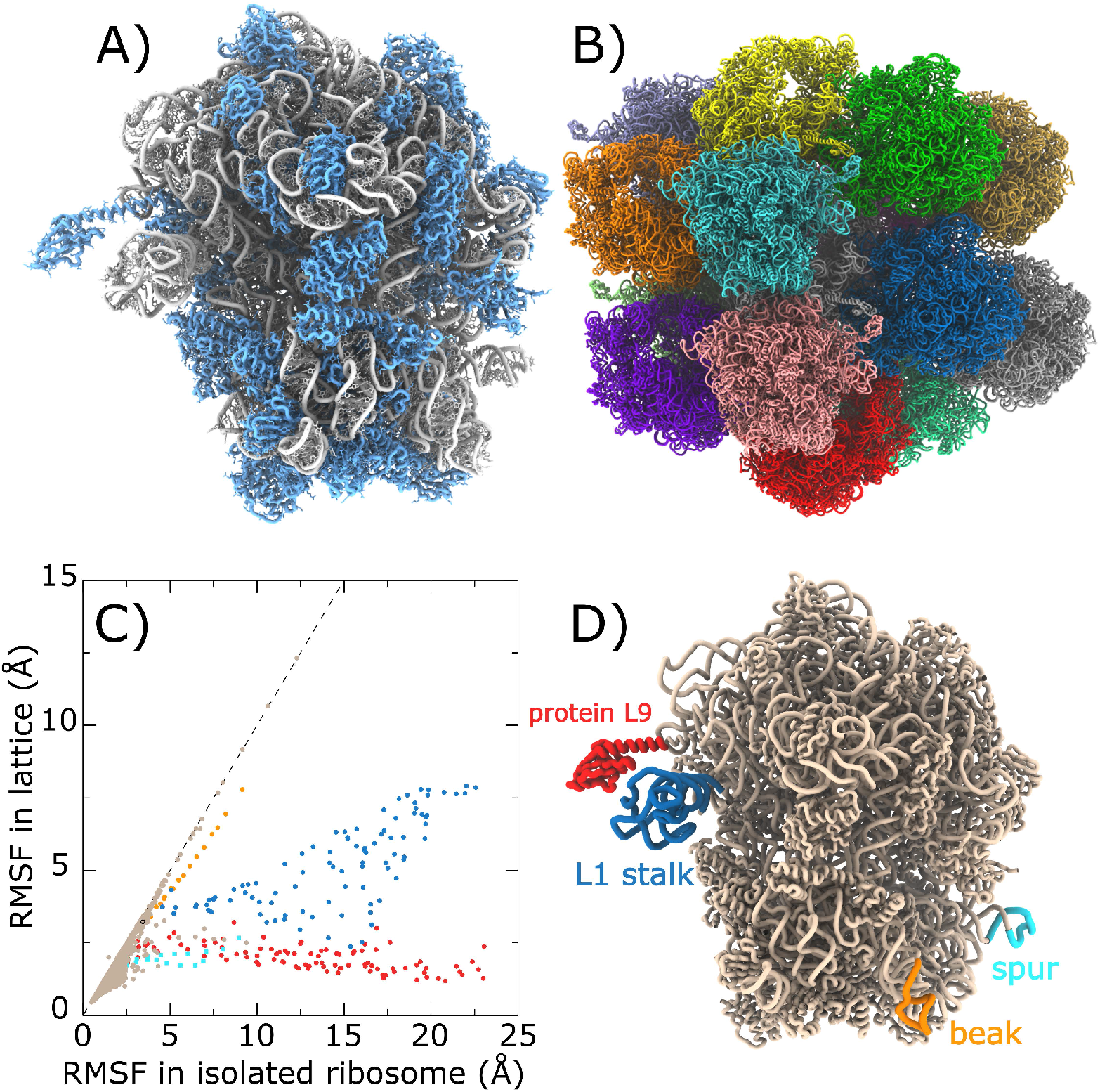
Probing the influence of crystallographic contacts on dynamics. A) Crystallographic structure of a bacterial ribosome (PDB ID:4XEJ^41^), with rRNA shown in white and proteins shown in blue. B) 15-ribosome cluster, where the arrangement corresponds to the experimental crystallographic lattice positions. The central ribosome (white) closely contacts 14 neighboring ribosomes (non-white). C) Spatial RMSF values, by residue, calculated from simulations of an isolated ribosome (panel A) and the central ribosome of the crystallographic cluster (panel B). Dashed line shows a slope of 1 (i.e. identical fluctuations in both systems). While the RMSF values of many residues are minimally impacted (light brown), there are multiple extended regions (protein L9, L1 stalk, 30S beak and spur) that are significantly more mobile in the isolated ribosome. D) Structure of the ribosome, colored by regions designated in panel C.

### Systems with many molecules

We next tested the ability of SMOG 2 to generate force fields for systems with large numbers of chains. For this test case, we generated a crystallographic lattice of a small, 64-residue protein. In the same manner as for the ribosome lattices described above, we generated crystallographic lattices of the protein CI2 (PDB ID: 2CI2^43^) that range from 1000 to 3000 unit cells. In the largest system, there were 36,000 proteins. All-atom SMOG models were successfully generated for all systems, again requiring approximately 11.2 minutes per million atoms (36k chains, 3.5 hours). To ensure the generated files were usable, each system was subsequently energy minimized and simulated at constant temperature using Gromacs. ^ii^

### Multi-million-atom chains

As a final stress test of the SMOG 2 software, we considered systems with extremely long chains. Here, we generated an artificial model of a nucleosome system. For this molecular model, we began with a single nucleosome structure (PDB ID 1P3I).^44^ We then extended the DNA strand using a model of an ideal linear B-form 40bp DNA that was generated with the W3DNA server.^45^ Using the Tk interface of VMD, ^46^ systems were generated with multiples of 100 nucleosome-linker segments. The largest system considered contained 1000 nucleosomes, with the dsDNA containing ≈ 184K basepairs and more than 3 million atoms per chain. This system also contained 8000 histone proteins, for a system total of 13,534,000 atoms that spans 4.5 *μ*m. Using a single compute core, SMOG v2.4 was able to generate an all-atom structure-based model for a chain of 1000 nucleosomes in ≈ 3.5 hours. For comparison, code limitations precluded the possibility of SMOG v2.3 to generate a force field for this system. As with the other cases, this system was energy minimized and simulated using Gromacs.^iii^

While this artificial system was used to demonstrate computational capabilities, it represents a notable milestone. That is, in modern models of chromatin, ^47^ the system is coarse grained at a resolution of 50000 base pairs per pseudoparticle. This extremely-coarse representation is suitable, since it corresponds to the resolution afforded by state-of-the art Hi-C measurements. With the presented advances in SMOG 2, the field may now consider a wide range of all-atom and coarse-grained (e.g. 1-3 beads per residue) models for systems that reach the scale of these experimentally-derived coarse-grained models of chromatin. This provides exciting opportunities, where SMOG models may be used to study the dynamics at “small” scales (i.e. 100,000 base pairs) and Hi-C-inspired models may be used to extend to the scale of full chromosomes. Together, this overlap of spatial scales can allow the field to establish a comprehensive understanding of chromatin dynamics that spans from atomic-level to chromosome-scale dynamics.

### Probing dynamics in subsystems

For biomolecular systems in which the dynamical phenomenon of interest involves a small sub-region of a larger assembly, it may be helpful to simulate only the dynamics of the sub-region, while approximating the dynamics at the boundary. With the SMOG 2 package, this can be achieved by extracting any subset of atoms from a larger system using the `smog_extract` tool. This tool also allows for position restraints to be imposed on boundary atoms (i.e. those that had interactions removed). It is possible to use isotropic^48^ or anisotropic^49^ restraints at the boundaries, such that the mobility of boundary atoms is consistent with the fluctuations observed in the full assembly. Below, we provide an example for how to use SMOG 2 to prepare a system and refine the boundary restraints.

To study a subsystem, we will use the `smog_extract` tool to construct a truncated ribosome system. In this example, an all-atom SMOG model was first generated for the full ribosome (Fig. 7A, PDB 6QNR50). Next, we extracted all atoms and interactions that are in the vicinity of the peptidyl transferase center (PTC) (Fig. 7B). Boundary atoms were automatically identified and assigned a default weight to each position restraint. Then, fol-lowing the methods of Noel et al.,^48^ the RMSF values of all boundary atoms were calculated and compared to the values found in the full ribosome. Based on this, the weight of each restraint was iteratively updated until the fluctuations of the boundary atoms were consistent with the full ribosome system (Fig. 7C). This yields a model for the subsystem in which artificial boundary effects are minimized. The utility of this strategy is that one may reduce the computational demand without unintentionally perturbing the dynamics due to the introduction of boundary restraints. In this specific system, one could then consider studying the interplay between ribosome flexibility near the PTC and the dynamics of the tRNA molecule.

**Figure 6:**
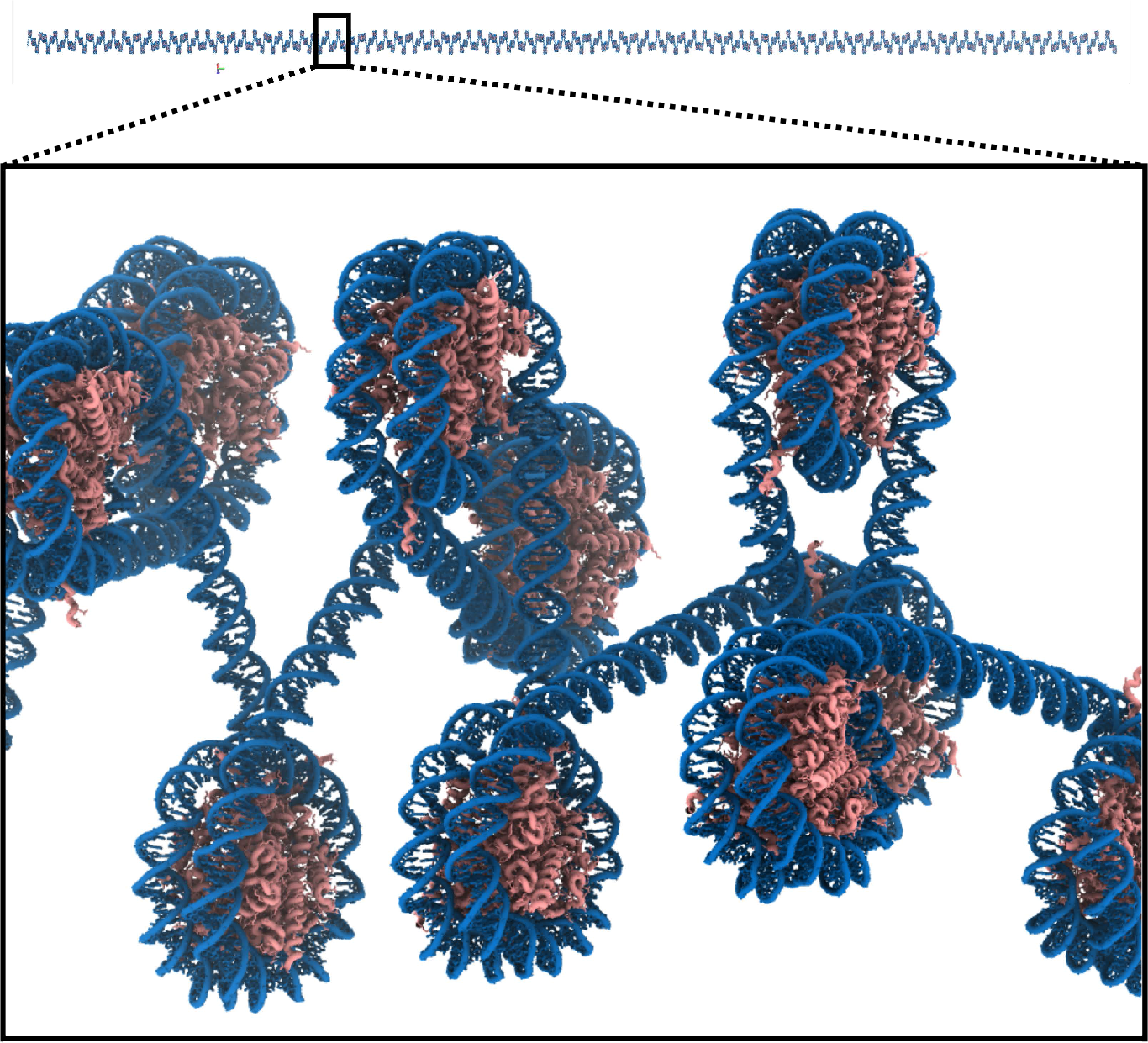
Simulating nucleosomes. top) Artificial 300-nucleosome system, where each DNA strand (blue) has 1131600 atoms. There are also 2400 histone proteins (pink), for a total of 4060200 atoms. bottom) Zoomed-in view of a 10 nucleosome segment that is associated with a single dsDNA helix. Recent improvements in SMOG 2 allow for the rapid generation of all-atom structure-based models for systems of this size. Systems that have up to 1000 nucleosomes were successfully tested.

**Figure 7:**
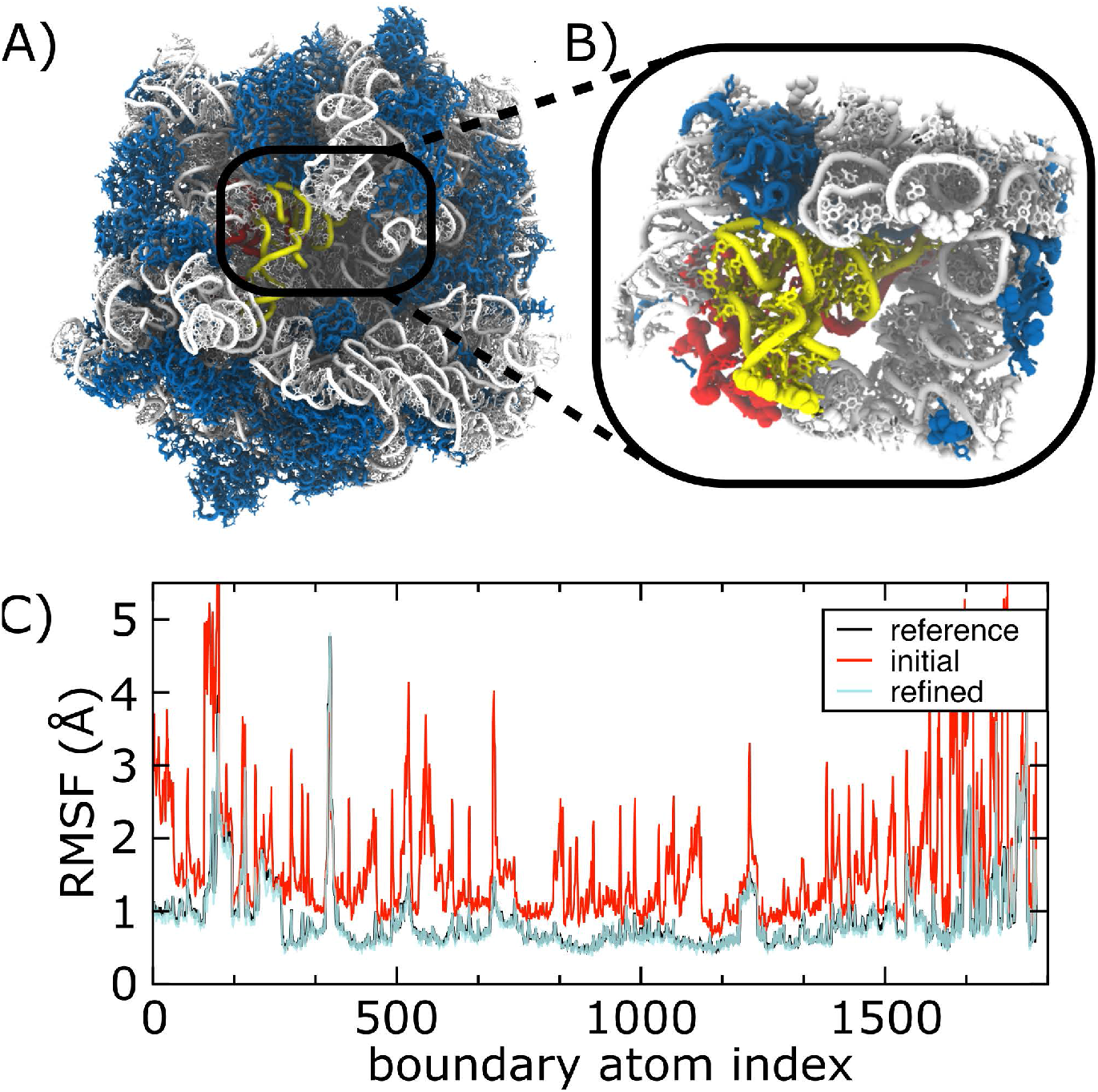
Simulating subsystems with SMOG. A) Structure of a complete bacterial ribosome (PDB 6QNR^50^), with the rRNA (white), proteins (blue) and tRNA (yellow and red) shown. B) After generating a structure-based model with SMOG 2, the `smog_extract` tool allows one to reduce the system size while preserving interactions present in the full assembly. Spheres indicate which atoms at the boundaries have interactions removed during truncation. Position restraints can be automatically applied to boundary atoms, to ensure the system maintains its overall structure. C) Position restraints may be refined,^48^ such that the dynamics of the subsystem boundary is consistent with the dynamics of the full assembly (labeled reference).

## Conclusions

SMOG 2 and OpenSMOG increase the accessibility of structure-based (SMOG) model development, application and dissemination. With the flexible construction of SMOG models that is afforded by SMOG2/OpenSMOG, these tools alleviate common technical challenges associated with force field development, while providing strategies that are tailored for the construction of structure-based models. By removing technical obstacles, SMOG2/OpenSMOG allows researchers to focus on posing deeper physical questions, which may be more easily answered through the systematic construction of SMOG model variants. In the current report, we highlight recent advances that include improved support for large molecular systems, as well as support for more elaborate SMOG model definitions. These representative applications illustrate only a few ways in which this platform can allow researchers to address broad biophysical challenges. In contrast to early structure-based models, these tools allow for the application of these models to systems/processes that span diverse length scales, ranging from the folding of small proteins to the dynamics of large-scale biomolecular assemblies.

## Author Information

### Author contributions

ABO, VGC, JKN and PCW wrote the software. All authors contributed to the design of the software. ABO, VGC, AH, SB, AW, YW, ED, JKN and PCW tested the software and performed simulations. All authors contributed to manuscript preparation.

### Notes

The authors declare no competing financial interest.

## Acknowledgement

PCW was supported by NSF grant MCB-1915843. This research was supported by the Center for Theoretical Biological Physics sponsored by the NSF (Grant PHY-2019745). JNO was also supported by the National Science Foundation (NSF) grants CHE-1614101 and PHY-1522550 and by the Welch Foundation (Grant C-1792). JNO is a Cancer Prevention and Research Institute of Texas (CPRIT) Scholar in Cancer Research. ABOJ acknowledges the Robert A. Welch Postdoctoral Fellow program. We would also like to acknowledge generous support from the Northeastern University Discovery cluster and Northeastern University Research Computing staff. We are also grateful for generous computational resources and support provided by the AMD COVID-19 HPC Fund program.

## Footnotes

i versions 5.1.4 and 2021.2 were both tested

ii versions 5.1.4 and 2021.2

iii versions 5.1.4 and 2021.2

